# The Microbial Ecology of Serpentinites

**DOI:** 10.1101/2024.11.10.622848

**Authors:** Daniel R. Colman, Alexis S. Templeton, John R. Spear, Eric S. Boyd

## Abstract

Serpentinization, the collective set of geochemical reactions initiated by the hydration of ultramafic rock, has occurred throughout Earth history and is inferred to occur on several planets and moons in our solar system. These reactions generate highly reducing conditions that can drive organic synthesis reactions potentially conducive to the emergence of life, while concomitantly generating fluids that are challenging for life owing to hyperalkalinity and limited inorganic carbon (and oxidant) availability. Consequently, serpentinite-hosted biospheres offer insights into the earliest life, the habitable limits for life, and the potential for life on other planets. However, the ability of serpentinites to support abundant microbial communities was only recognized ∼20 years ago with the discovery of deep-sea hydrothermal vents emanating serpentinized fluids. Here, we review the microbial ecology of marine and continental serpentinite-hosted biospheres in conjunction with a comparison of publicly available metagenomic sequence data from these communities to provide a global perspective of serpentinite ecology. Synthesis of observations across global systems reveal consistent themes in the diversity, ecology, and functioning of communities. Nevertheless, individual systems exhibit nuances due to local geology, hydrology, and input of oxidized, near-surface/seawater fluids. Further, several new (and old) questions remain including the provenance of carbon to support biomass synthesis, the physical and chemical limits of life in serpentinites, the mode and tempo of *in situ* evolution, and the extent to which modern serpentinites serve as analogs for those on early Earth. These topics are explored herein from a microbial perspective to outline key knowledge-gaps that should be a focus of future research.

## INTRODUCTION

The earliest life was supported by chemical energy that was released when disequilibria in reduced and oxidized chemicals (electron donor and acceptors) were dissipated (1). While uncertainty remains in the timing of the emergence of photosynthesis (e.g., between ∼4 and 3 Ga (2)), life dependent on chemical energy is widely recognized to have predated that dependent on light energy (1, 2). Mantel convection, volcanism, and water-gas-mineral interactions were critical in generating chemical disequilibria among electron donor/acceptor pairs to support chemosynthetic life during this time. Of particular significance is the process of serpentinization, or the collective series of chemical reactions that occur upon the hydration of (ultra)mafic rock (iron-rich, silica-poor rock) resulting in the oxidation of iron-bearing minerals and the formation of serpentine [(MgFe(II)Fe(III))_3_ (Fe(III)Si)_2_O_5_(OH)_4_]. Serpentine formation is accompanied by hydrogen gas (H_2_) generation (3) that, when accumulated to high enough concentrations, can drive the reduction of inorganic carbon, leading to the production of carbon monoxide (CO), formate (CHO_2_^-^), methane (CH_4_) (3–5), and even longer chain compounds like acetate (C_2_H_3_O_2_^-^) and pyruvate (C_3_H_3_O_3_^-^) (6).

Perhaps unsurprisingly, many origin of life theories invoke the geological process of serpentinization and a primitive autotrophic energy metabolism founded on H_2_, e.g., the Wood-Ljungdahl (WL) pathway of carbon dioxide (CO_2_) fixation (7–10). The WL pathway is thus far the only known carbon fixation pathway present in both Archaea and Bacteria and is the least energy intensive (11). Further, many hydrogenotrophic acetogenic Bacteria, methanogenic Archaea, and sulfate (SO_4_^2-^) reducing Bacteria/Archaea, among other anaerobes, remain dependent on the WL pathway as part of their core energy-conserving and CO_2_-fixing pathways (12). Thus, serpentinization and microbial physiologies theorized to be dependent upon serpentinization by-products have remained a major focus of early life studies.

(Ultra)mafic rock like peridotite that primarily comprises olivine [e.g., (Mg, Fe)2SiO4] and pyroxene [e.g., orthopyroxene: (Mg,Fe)_2_Si_2_O_6_]) is common in the Earth’s mantle and tectonically uplifted regions of the oceanic crust, but is less common on continents, except where these rock units are obducted onto the crust, in what is termed an ophiolite. While the geochemistry, mineralogy, and geology of serpentinites have been widely studied for many decades, their potential to host microbial communities remained essentially unknown until only ∼20 years ago when H_2_-enriched fluids from deep-sea hydrothermal vents in the Lost City Hydrothermal Field (LCHF; Atlantis Massif) were shown to host abundant microbial communities (13). Since this discovery, interest in serpentinite-hosted microbial ecosystems has grown considerably and now includes studies aimed at understanding the extent and nature of Earth’s subsurface biospheres (14, 15), the earliest life on Earth (7, 9, 16), the potential for life on other planetary systems (17), the environmental limits to life (18), and most recently, the potential for geological H_2_ production and CO_2_ sequestration (19, 20). This has included investigations of a variety of serpentinites, including marine systems at the LCHF, Prony Bay Hydrothermal Field (PBHF; New Caledonia), and the Old City Hydrothermal Field (OCHF; Southwest Indian Ridge), as well as terrestrial ophiolites in California, USA (at the Cedars Springs, Ney Springs, and the Coast Range Ophiolite Microbial Observatory; CROMO), the Samail Ophiolite (Oman), and the Voltri Massif (Italy), among others (13, 21–27).

Collectively, these studies have revealed the compositions and functions of microbial communities that inhabit these chemically diverse environments and have provided insight into both the abiotic and biotic factors that influence their ecological functioning. Despite the considerable insights generated over the past ∼20 years, an equal number of fundamental new questions have arisen. Here, we review insights into the microbial and evolutionary ecology from the past two decades of microbial investigations and outline key unresolved questions to help move serpentinite studies forward. To further this effort, publicly available metagenomic sequence data were compiled from both marine and terrestrial serpentinite-hosted microbial communities and were subjected to comparative analyses.

## 1. TAXONOMIC AND FUNCTIONAL DIVERSITY

### 1.1 Methanogens in Serpentinite Communities

The discovery of serpentinizing systems as microbial habitats coincided with a renewed interest in subsurface environments as globally significant reservoirs of microbial diversity and biomass (28–30). Key to sustaining these subsurface microbial ecosystems is an abundance of geogenic reductant (e.g., H_2_) that can be coupled with inorganic oxidants (e.g., CO_2_ and SO_4_^2-^) to conserve energy that then fuels primary production (30–32). Thus, lithoautotrophic microorganisms in serpentinization-influenced systems became prime targets for investigation, including, in particular, archaeal hydrogenotrophic methanogens that couple H_2_ oxidation to CH_4_ production (CO_2_ + 4H_2_ → CH_4_ + 2H_2_O) and bacterial (homo)acetogens that couple H_2_ oxidation to acetate production (4H_2_ + 2CO_2_ → CH_3_COOH + 2H_2_O) (16), among other microbial metabolic guilds. Indeed, early microbiological studies of the LCHF revealed dominance by methanogenic Archaea, and particularly by a single 16S rRNA gene phylotype referred to as the Lost City Methanosarcinales (LCMS) in vents that are highly impacted by serpentinization (33, 34). Analyses from later metagenomic studies indicated that the LCMS are potentially capable of hydrogenotrophic methanogenesis in addition to acetoclastic (acetate fermentation) and/or methylotrophic (methyl-group fermentation) methanogenesis (35, 36). Microcosm carbon stable isotope studies also showed a stimulation of CH_4_ production when amended with H_2_, and surprisingly, also suggested the capacity for anaerobic methanotrophy (37). However, considerable debate remains whether canonical methanogens can also conduct anaerobic methanotrophy (38) and further physiological and biochemical study of these uncultured taxa is still needed. Nevertheless, these early studies primarily investigated biofilm communities within vent chimney interiors, potentially limiting the perspective of taxonomic and functional diversity associated with serpentinizing systems.

More recent studies of LCHF waters have documented greater variability in community compositions, including those that are dominated by bacterial taxa, with LCMS generally comprising subdominant populations (35). Similarly, studies of other serpentinite systems have documented a variable presence and abundance of methanogens, ranging from systems like the Tablelands (Canada), CROMO, Ney Springs, or Hakkuba Springs (Japan) where they were not detected (24, 39–41) to systems like the Samail Ophiolite or LCHF where they are present or abundant but only in some communities (35, 42–44). Yet in other systems, such as springs of the Santa Elena Ophiolite (Costa Rica) methanogens can be dominant (45). Consistent with these studies, an assembly-free taxonomic analysis of 70 publicly available metagenomes (**Fig. 1**; see Supplementary Information for methods) from serpentinite systems suggests a paucity of methanogenic taxa in most systems, although considerable variation in their distribution and abundance exists within and among communities, particularly from the Samail Ophiolite and LCHF (**Fig. 2**). Thus, while methanogens and methanogenesis have been a primary focus of microbiological studies in serpentinite systems since their initial discovery in the LCHF, additional studies of serpentinites over the last ∼20 years have reshaped our overall understanding of their microbial ecology. Collectively, these findings prompt the fundamental question of what environmental or biological factors promote or constrain the distribution, abundance, and metabolism of methanogens in serpentinites?

**Fig. 1.**
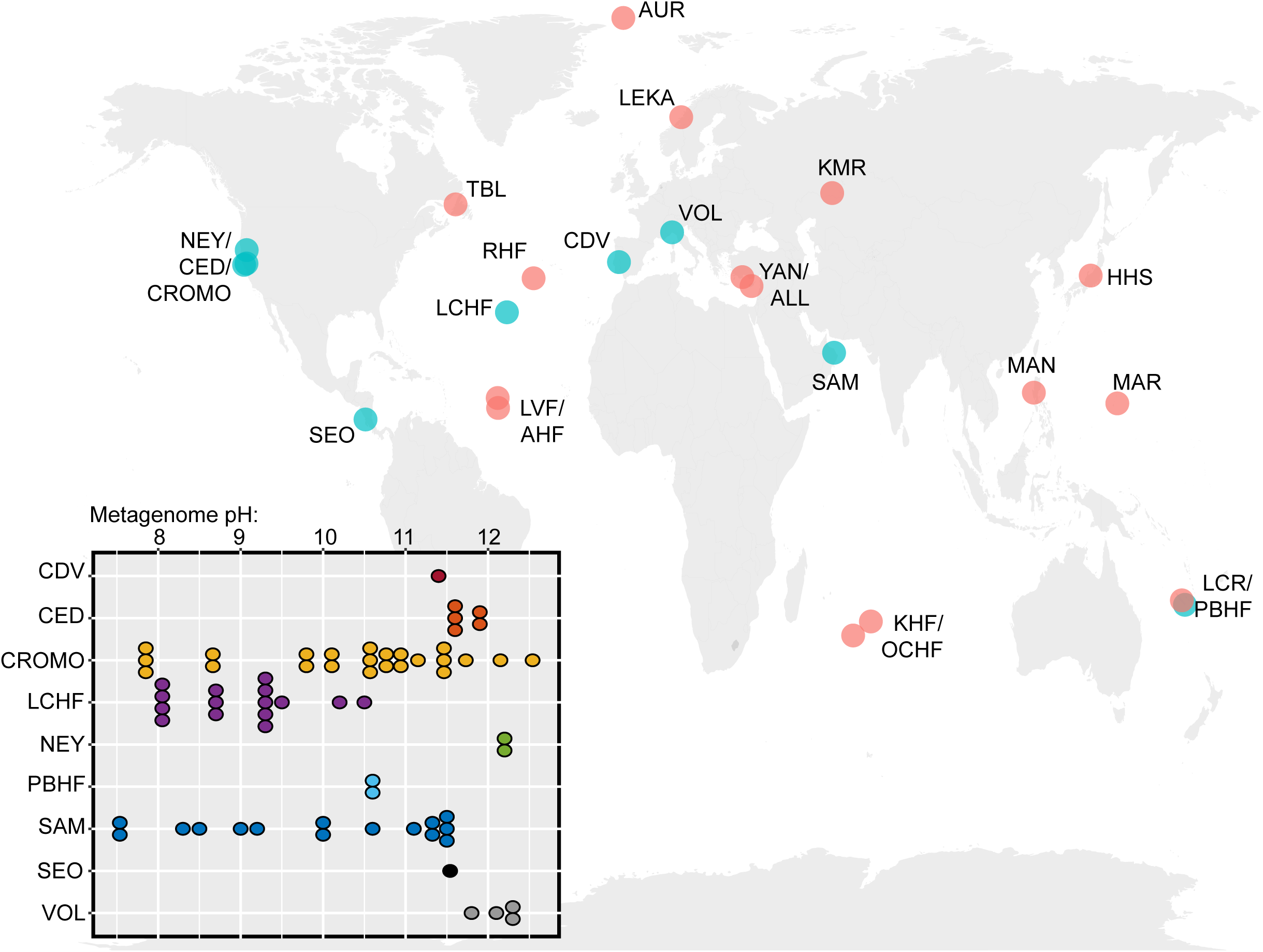
Global distribution of serpentinization-influenced environments that have been investigated microbiologically and those that hosted 70 publicly available shotgun metagenomes used in the meta-analysis of this review. Geologic systems are indicated by points and abbreviations on the map, with those in green hosting metagenomes that were used in the meta-analysis of this review. Bottom left insert shows the pH distribution of water samples from which the metagenomes were derived, as reported by the original publication. Additional information for metagenomes is shown in Supplementary **Table 1**. Abbreviations are defined as follows (and further identified in Supplementary **Table 1**). NEY: Ney Springs; CED: The Cedars; CROMO: Coast Range Ophiolite Microbial Observatory; SEO: Santa Elena Ophiolite; TBL: Tablelands Ophiolite; RHF: Rainbow Hydrothermal Field; LCHF: Lost City Hydrothermal Field; LVF: Logatchev Vent Field; AVF: Ashadze Vent Field; AUR: Aurora Seamount; LEKA: Leka Ophiolite; CDV: Cabeço de Vide; VOL: Voltri Massif; YAN: Yanartaş; ALL: Allas Springs; SAM: Samail Ophiolite; KMR: Khalilovsky Massif; KHF: Karei Hydrothermal Field; OCHF: Old City Hydrothermal Field; HHS: Hakuba Happo Hot Springs; MAN: Manleluag Spring; LCR: La Crouen Spring; PBHF: Prony Bay Hydrothermal Field.

**Fig. 2.**
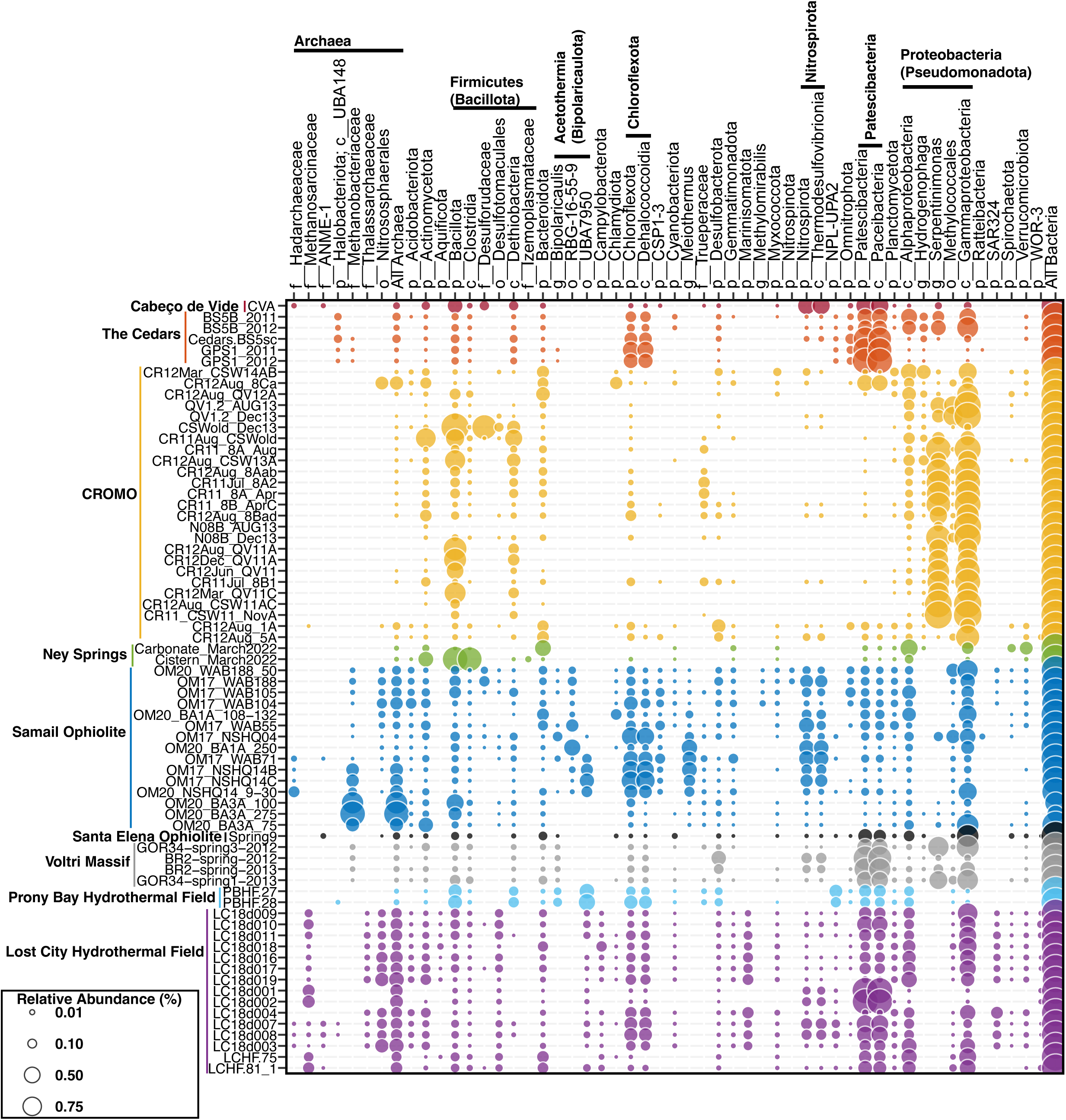
Taxonomic composition of 70 community metagenomes from serpentinization-influenced environments. Each row represents a metagenome organized by site and by increasing pH from top to bottom (within each site). Each column represents a taxonomic group that was identified in >1% relative abundance in at least one of the metagenomes. Metagenomic data and publicly available accession information is provided in **Supplementary Table 1**. Community composition was evaluated from raw reads using the SingleM pipeline (described in Supplementary Methods). Taxonomic groups are defined as in the Genome Tree Database (GTDB) at various taxonomic levels: p: phylum; c: class; o: order; f: family; g: genus. Sub- phylum level taxa that are discussed in the manuscript are specifically included. Circles are colored by their corresponding system and sized according to their estimated relative abundance, as indicated by the legend on the bottom left.

### 1.2 A Revised Understanding of Methanogens and other Archaea in Serpentinite Ecosystems

Members of the LCMS (family Methanosarcinaceae) have been identified in other globally distributed serpentinite systems, including in the PBHF (46), while a LCMS-like phylotype (TCMS) has been observed in the Cedars (47). Thus, LCMS-like methanogens appear to mostly be restricted to marine serpentinite systems. In contrast, *Methanobacterium*-related methanogens (class Methanobacteria) are commonly identified in terrestrial serpentinites, including at the Cedars, Ney Springs, Samail Ophiolite, Zambales (Philippines), and Voltri Massif systems (27, 42, 43, 48–52). Like the LCMS in marine systems, the *Methanobacterium*- related methanogens found in terrestrial systems comprise only a single or a few genomic phylotypes (48, 53), suggesting specific adaptations that facilitate their competitiveness in these environments.

Serpentinite-associated *Methanobacterium* primarily conduct hydrogenotrophic methanogenesis based on cultivated representatives (51, 54), genomic inference (48, 50, 53), or ^14^C carbon isotope uptake experiments (48), consistent with most cultured Methanobacteria (55). However, several unique adaptations have been identified among Methanobacteria in serpentinites, including hypothesized formate-driven (rather than H_2_-driven) methanogenesis in *Methanobacterium* spp., as revealed by metagenomics and isotopic studies in communities from the Samail Ophiolite (48, 56) (discussed in greater detail in **Section 3.2**). Further, the potential for formate-dependent methanogenesis by *Methanobacteria* in springs of the Voltri Massif was also suggested by metagenomics and ^13^C carbon substrate conversion studies (27). Notably, spring waters from the Santa Elena Ophiolite harbored 16S rRNA gene evidence for many of the above taxa, in addition to others not commonly identified in serpentinizing systems (Methanomicrobiales), that were variously implicated in H_2_-, formate-dependent, or other types of methanogenesis (45). In addition to methanogenic Archaea, anaerobic methanotrophic Euryarchaeota (ANME) were identified in the LCHF (33), and more recently in the Santa Elena, Cabeco de Vide, and Samail Ophiolite systems (43, 45, 57). The ANME have been hypothesized to participate in syntrophic anaerobic oxidation of CH_4_, which is often abundant in fluids impacted by serpentinization (58). Thus, the distribution of both methanogens and ANME in serpentinite systems is patchy and restricted to only a few archaeal phylotypes within a few divisions. This conclusion is consistent with the assembly-free analysis of 70 serpentinite metagenomes conducted herein, which also reveals a patchy distribution of methanogenic or anaerobic methanotrophic archaeal taxa (**Fig. 2**).

Beyond the relative paucity of methanogenic or ANME archaeal phylotypes identified in serpentinizing systems, few other archaeal taxa have been identified (**Fig. 2**). These include representatives of Nitrososphaeria (21, 22, 59) that comprise ammonia-oxidizers typical of fresh- and sea-waters (60) and *Thalassoarchaea* typical of global marine waters (61) that have been detected in marine serpentinite communities (21, 46). Likewise, anaerobic Hadesarchaeota putatively capable of heterotrophic metabolism that is supplemented by CO oxidation (50) and that are common in subsurface environments (62) were recently identified in highly reduced fracture waters of the Samail Ophiolite (63) and in Zambales springs (50), but may also be present as low-abundance taxa in other systems (**Fig. 2**). Collectively, while these data reveal that Archaea can inhabit serpentinites, they tend to be rare, only comprise a few total phylotypes, and are generally subdominant to Bacteria. Intriguingly, serpentinites are generally characterized as hosting waters that are highly reduced, nutrient limited, anoxic, and generally energy limited, conditions that tend to favor archaeal inhabitation (64). Thus, the general lack of archaeal diversity or abundance in these systems remains perplexing.

### 1.3 Recent Discovery of Acetogen-like Bacteria as Putative Primary Producers

In contrast to the Archaea, diverse bacterial taxa with equally diverse metabolic capacities have been identified in serpentinite systems. Anaerobic bacteria are particularly prevalent in serpentinized waters, including those involved in sulfur cycling (primarily sulfate reduction) and those putatively capable of acetogenesis or 1-carbon metabolism. Evidence for the presence of bacterial acetogens has been much less forthcoming compared to that for methanogens, with no acetogenic isolates yet reported from serpentinization habitats. Rather, genome-resolved metagenomics approaches (e.g., via metagenome assembled genomes, MAGs) have only recently provided evidence for several putatively acetogen-like organisms from thus far uncultured bacterial divisions (35, 39, 65–67). These putative acetogen MAGs represent populations that are often dominant members of, and putative primary producers in, serpentinized terrestrial and marine communities (35, 46, 65). Further, while heterotrophic acetogenesis (acetate production from organic compounds) has been demonstrated in Cedars communities using carbon isotopic microcosm experiments (68), evidence for autotrophic acetogenesis (homoacetogenesis) has yet to be demonstrated or evaluated in a serpentinizing system with experimental approaches. Canonical homoacetogenesis is generally considered to be a challenging metabolism to support life owing to the minimal free energy released from the H_2_/CO_2_ redox couple (4H_2_ + 2CO_2_ → CH_3_COOH + 2H_2_O couple; ΔG = –40 kJ per mole under physiological conditions) (69). However, the thermodynamics of this reaction become especially problematic in highly serpentinized fluids (39) and are exacerbated by the limited availability of dissolved inorganic carbon (DIC) in these systems (44). Autotrophic acetogenesis may consequently require metabolic adaptations to serpentinite environments that help overcome these challenges, such as the use of formate rather than CO_2_ to drive acetogenesis and recruitment of electron bifurcating enzyme systems to drive exergonic electron transfers (39, 65, 70), as discussed in greater detail in **Section 3.2**. Intriguingly, a recent genome-resolved metagenomic analysis identified an archaeon within one of the 7 canonical groups of methanogens (Methanocellales) that was abundant in springs of the Cedars, but that appear to have relatively recently lost the capacity for methanogenesis and are inferred to be non-H_2_- dependent acetogens (71). These results are consistent with laboratory experiments demonstrating that canonical methanogens can be engineered to conduct acetogenesis via targeted disruptions to methanogenesis pathways (72). Although further confirmation of these recent results are needed, the putatively acetogenic Methanocellales represent a previously unrecognized lineage of acetogens and confirmation of a novel evolutionary scenario whereby methanogenic Archaea diverge into non-methanogenic acetogens. Acetate is often one of the most abundant short chain carbon compounds in serpentinite waters (42), suggesting the need for additional studies aimed at identifying its biotic or abiotic source and fate in these ecosystems.

### 1.4 Other Prevalent Bacterial Functional Guilds

Sulfur-metabolizing bacteria, and particularly sulfate-reducing bacteria (SRBs), are commonly abundant in serpentinite-hosted communities, including taxa related to the *Thermodesulfovibrio* (class Thermodesulfovibrionia) (26, 35, 59), *Dethiobacter* (class Dethiobacteria), *Desulforudis* (family Desulforudaceae), *Desulfotomaculum* (order Desulfotomaculales), and members of the Desulfobacterota class (previously Deltaproteobacteria) (35, 42, 73), among others. Further, rates of sulfide production coordinated with H_2_ drawdown, ^13^C substrate uptake experiments, and molecular microbial analyses have suggested that SRB may be important primary producers and consumers of H_2_ in the LCHF system (74, 75) and prevalent in the subsurface of serpentinizing systems, as in the Mariana Forearc based on sulfur isotope fractionation analysis of fluids (76). Moreover, ^35^S radiotracer microcosm uptake studies suggest that SRB from the Samail and Coast Range Ophiolite Microbial Observatory (CROMO) systems are active across the range of pH (e.g., pH ∼7 - 12.5) encountered there, although their activity may be limited at higher pH (77).

As in most subsurface environments, SO_4_^2-^ is one of the most abundant oxidants for lithotrophic metabolisms in serpentinizing systems, either due to input of seawater with high SO_4_^2-^ (∼28 mM) or to radiolytic- or radical-based abiotic pyrite oxidation (78, 79). The thermodynamics of SO_4_^2-^ reduction with H_2_, CO, or organic acids (e.g., acetate or formate) is more favorable than CO_2_ reduction through methanogenesis or acetogenesis (74) partially explaining why SRB often outcompete other metabolic guilds in many anaerobic natural systems (80). SRB have also been traditionally considered to outcompete methanogens for H_2_ in freshwater environments when SO_4_^2-^ >30 μM (81). Consistent with these assertions, community composition-informed niche modeling of hyperalkaline spring waters of the Samail Ophiolite and elsewhere suggest *Thermodesulfovibrio* can outcompete *Methanobacterium* when SO_4_^2-^ concentrations exceed 10 μM (82). However, the co-occurring potential for SO_4_ ^2-^ reduction and methanogenesis as assessed via microcosm studies and genomic analyses, for example in Samail ophiolite subsurface waters (42, 77), suggests that their niches are temporally and/or spatially defined by parameters other than oxidant availability alone, thereby minimizing niche overlap and enabling co-existence.

Less well understood in serpentinite systems are the roles of microorganisms in the oxidative sulfur cycle (e.g., in metabolizing S_2_O_3_^-^, S^2-^, or S^0^), although 16S rRNA gene and genomic investigations have suggested a high potential for sulfur oxidation in several systems (24, 73, 83). Further, aerobic S_2_O_3_^-^ oxidizing (Rhodobacteraceae and *Halomonas* spp.) and sulfur-compound disproportionating (*Dethiobacter* sp.) strains have been isolated from the Ney Springs and CROMO systems (24, 84). Oxidative sulfur cycling metabolisms require the availability of oxidants (e.g., O_2_, or NO_3_^-^) with which to couple to sulfur-compound oxidation. Consequently, these environments are likely to be largely restricted to near surface zones of terrestrial serpentinites or near the sea water/serpentinized fluid interface in marine systems (24, 83).

Conspicuously missing from most surveys of microbial taxonomic and functional diversity in serpentinizing systems is evidence for Fe(III)-reducing or Fe(II)-oxidizing populations, given the abundance of Fe-bearing minerals in host-rock settings and the critical role that these minerals play in serpentinization reactions. Fe(III) reducing enrichment and isolate cultures were recovered from the hyperalkaline Allas Spring (Cyprus) (85), in addition to a magnetite-reducing *Paenibacillus* sp. (Firmicutes phylum) from the Cedars systems (86). It is worth noting that the metabolic pathways of dissimilatory Fe(III) reduction or Fe(II) oxidation are not conserved, can be non-specific, and remain partially unknown (87), thereby confounding inference from genomic-based investigations. Consistent with this perspective, the genus *Dethiobacter* was originally described via moderately alkaliphilic strains (e.g., *D. alkaliphilus*) isolated from soda lake sediments that were capable of H_2_-based reduction of various sulfur compounds (e.g., S_2_O_3_^-^ or polysulfides) (88). *Dethiobacter* spp. are common inhabitants of serpentinization-influenced waters, as indicated above (and in **Fig. 2**), and *D. alkaliphilus* has recently been shown to comprise strains that reduce Fe(III) with various electron donors including H_2_ and organic acids, in addition to strains that can putatively oxidize Fe(II) anaerobically (89). Given that phylotypes closely related to these strains are ubiquitous in serpentinite environments (89), it is possible that they are conducting similar metabolisms in these systems. Consistent with this supposition, a previous revisiting of several described alkaliphilic, sulfur-reducing taxa revealed the widespread ability to reduce Fe(III) via multiheme cytochrome *c* proteins or other non-specific pathways (90). Consequently, dissimilatory Fe-based metabolisms may be more prevalent in serpentinite environments than previously recognized due to difficulties in identifying these activities from culture-independent analyses.

Numerous bacterial taxa capable of aerobic respiration have also been identified in serpentinization-influenced waters (18, 47, 63, 91). Although waters in serpentinites are generally considered to be highly reducing and anoxic, mixing of these fluids with near surface oxygenated waters or the infusion of atmospheric gases at spring/wellhead air-water interfaces provides substantial opportunity for microbial metabolisms to leverage surface-derived oxidants (e.g., O_2_). Often, these aerobes couple with reductants generated from serpentinization (e.g., H_2_ or CH_4_) in highly exergonic reactions (92). For example, the genus *Serpentinimonas* (class Betaproteobacteria) was isolated as an H_2_-dependent aerobic autotroph from Cedars spring waters (18), and exhibited optimal growth at pH of ∼11 (with growth up to pH 12.5). Molecular analyses of serpentinized fluids from terrestrial serpentinites commonly identify this genus (41, 47, 93), as also observed in the assembly-free analysis conducted here (**Fig. 2**).

Other diverse aerobic taxa are commonly identified in surface-exposed continental systems (24, 27, 49) and marine vent fluids (22, 34), suggesting a widespread potential to leverage the dynamic niche space at the interface of reduced serpentinized fluids and oxidized surface/sea waters. In addition, Bacteria inferred to be obligate aerobes are commonly found in deep waters isolated from near surface inputs, including for example, *Meiothermus*, whose abundances paradoxically increased with depth in a highly reducing well within the Samail Ophiolite (63). An intriguing, but so-far little explored, alternative explanation for apparent O_2_ availability in anoxic serpentinite system waters may occur through “dark” O_2_ production via several non-photosynthetic mechanisms that have only recently been identified as ubiquitous among microorganisms. This includes O_2_ production through dismutation of chlorite (ClO_2_^-^ →Cl^-^ + O_2_), nitric oxide (2NO → N_2_ + O_2_), and hydrogen peroxide (2H_2_O_2_ → 2H_2_O + O_2_), as well as during oxidation of ammonia via an unknown mechanism (94, 95). Such reactions have recently been inferred to support many subsurface ecosystems that have long been isolated from atmospheric sources of O_2_ (96). While genomic evidence for NO and H_2_O_2_ dismutation in serpentinite communities is limited, genes encoding chlorite dismutase (Cld) proteins are widespread and oftentimes abundant, in particular in studies of subsurface waters within the Samail Ophiolite (63). Attempts to detect ClO_2_^-^ (or perchlorate that can yield ClO_2_^-^ when reduced) in Samail Ophiolite fracture waters have thus far been unsuccessful (63), which may point to its rapid biological consumption. Additional research is needed to assess the potential for “dark” O_2_ production in supporting subsurface biospheres in serpentinites and to assess potential sources of ClO_2_^-^, NO, and H_2_O_2_ in these systems.

## 2. ADAPTATIONS TO OVERCOME PHYSIOLOGICAL CHALLENGES IN SERPENTINITE ECOSYSTEMS

### 2.1 Challenges for Life in Highly Serpentinization-Influenced Environments

Despite the diversity of microbial taxa supported by chemical products of serpentinization (e.g., H_2_, HCOO^-^, and CH_4_), the suite of geochemical reactions responsible for the formation of these products also present distinct challenges for microbial life. In particular, mineral replacement reactions during water-rock interactions in serpentinites can lead to alkaline and hyperalkaline pH (e.g., >12 (97)), dissolved inorganic carbon (DIC) concentrations that are critically low, often below detection (10uM) (13, 42, 59, 84), highly reduced waters/minerals (e.g., redox potentials as low as -750 mV (26, 59, 98), and limited availability of oxidants that can be used to support microbial metabolism. Consequently, while highly serpentinized waters may have abundantly available low potential reductant to fuel metabolism (e.g., of H_2_ or CH_4_; often in millimolar concentrations) (42, 74, 84), the coupling of such reductants to oxidants that are at low concentrations can be problematic. This phenomenon, combined with physico- chemical stresses like hyperalkaline pH and extremely low redox potentials, impose substantial bioenergetic burdens on microbial cells (15). The combination of extreme conditions and the stress that it imposes is likely responsible for the low cell abundances in serpentinite fluids, with cell concentrations ranging as low as 10^2^ - 10^3^ cells ml^-1^ (27, 47) and up to 10^4^-10^5^ cells ml^-1^ (27, 35, 45, 52).

Because the pH of fluids increases during the process of serpentinization, pH itself is often used a proxy for the progress of the collective reactions involved in serpentinization. Waters minimally impacted by serpentinization exhibit circumneutral pH while those more impacted by serpentinization can have pH as high as >12 (97). However, all known microbial cells maintain intracellular pH near ∼7, with few exceptions (99, 100). Consequently, maintaining near-neutral intracellular pH in the face of several orders of magnitude difference in extracellular H^+^/OH^-^ availability is a key challenge to overcome for cells inhabiting highly serpentinized waters. A common adaptation in alkaliphiles for maintaining cellular ionic homeostasis is the cross-membrane transport of ions, for instance by antiporters that can exchange intracellular Na^+^ or K^+^ for scarce extracellular H^+^, as exemplified by the multiple resistance and pH adaptation (Mrp) antiporter system (100). Consistently, genes encoding Mrp antiporters (in addition to those that can exchange Ca^2+^, K^+^, and other cations) are commonly enriched in the genomes of various serpentinite-associated taxa identified by metagenomics- (39, 50, 66) or cultivation-based analyses (101). This implies an important role for antiporters in maintaining intracellular pH homeostasis in hyperalkaline waters generated by serpentinization.

Numerous other cellular-level adaptations to mitigate the loss of intracellular H^+^ and promote the influx of extracellular H^+^ have been identified in alkaliphiles, including cell wall structural adaptations that repel anions, the use of secondary cell wall polymers that can trap H^+^, plasma membrane fatty acid compositions that moderate OH^-^/H^+^ exchange, and the up-regulation of metabolic activities that produce organic acids (e.g., lactic acid) to moderate intracellular pH, among others (as reviewed in (100, 102). Undoubtedly, combinations of these adaptations are also commonly employed by serpentinite-hosted microbial taxa, although relatively few isolates have been recovered from these systems (91) and many of these adaptations lack known genomic markers that can be used to infer their presence in the absence of cultivation-based studies. A membrane-associated protein that has received less attention for its potential to enhance the fitness of microorganisms inhabiting hyperalkaline serpentinite environments are extracellularly- oriented [NiFe]-hydrogenases (typically identified genomically via a signal peptide on the N-terminus of their large subunit) that are used to generate reducing equivalents for microbial metabolism, including in some archaeal methanogens and bacterial sulfate reducers (103, 104). H_2_ oxidation activity outside of the membrane (e.g., in the periplasmic space of sulfate reducers) leads to a buildup of protons that can be used to conserve energy. Of potential equal importance is a role in the generation of localized acidity through [NiFe]-hydrogenase-mediated H_2_ oxidation that could enhance the dissolution of carbonates (and other minerals), thereby liberating dissolved inorganic carbon (DIC) to overcome DIC limitation (discussed in more detail in section **3.2**). Indicators of carbon assimilation activity have been shown to outweigh energy conservation activity in communities inhabiting the most serpentinized waters in Oman (52). Thus, future studies investigating autotrophic isolates that encode predicted extracellular- oriented, membrane-bound [NiFe]-hydrogenases could be used to more definitively probe the potential for this adaptative strategy in hyperalkaline, DIC-poor systems.

In addition to the need to maintain a near-neutral intracellular pH, nearly all life relies on cross-membrane chemiosmotic potential gradients to conserve energy and power ATP synthesis (105, 106), wherein H^+^ gradients are used in most organisms (107). Specifically, elevated extracellular H^+^ concentrations are established to form a cross-membrane gradient that can drive ATP synthesis via one of three ATP synthase variants (F-, and V-, and A-Types) (108). However, alkaliphiles can use Na^+^ gradients (109) that may be established by the antiporter activities described above and that can be utilized by ATP synthase variants structurally adapted to specifically exchange Na^+^ (108). Consequently, the use of Na^+^ chemiosmotic potentials may be an adaptation for cells that inhabit the most hypersaline serpentinized fluids.

Many taxa that inhabit low-energy anoxic environments (e.g., fermentative and acetogenic bacteria) were historically presumed to persist at the thermodynamic limits of life by synthesizing ATP via only substrate level phosphorylation (SLP). However, the relatively recent discovery of chemiosmotic potentials (predominantly Na^+^-driven) in autotrophic acetogens via the membrane-associated *Rhodobacter* nitrogen fixation (Rnf) complex (69) has helped resolve the energetic basis for many anaerobic metabolisms, in conjunction with other recently discovered metabolic strategies (e.g., electron bifurcation; (105)). Rnf couples the cytoplasmic oxidation of reduced iron-sulfur proteins (ferrodoxins) to NAD^+^ reduction and cross membrane Na^+^ translocation, allowing the development of a chemiosmotic potential generation under H^+^- limiting conditions. Rnf has been recently identified in the genomes of putative acetogenic bacteria in hyperalkaline fluids of the Samail Ophiolite and the Cedars (39, 65, 66), in addition to numerous MAGs from LCHF fluids (35) and cultivars from the Cedars (101). Thus, the use of Rnf complexes may represent an additional energy conservation adaptation to hyperalkaline conditions in serpentinites. Nevertheless, among the few isolates recovered from serpentinizing systems, members of the bacterial genus *Serpentinimonas* do not exhibit preferential use of Na^+^ to drive ATP synthesis via chemiosmotic potential (18), nor do three other alkaliphilic isolates recovered from the Cedars (101). Nevertheless, much remains unknown about the selectivity of ATP synthases for H^+^ or Na^+^ (108), with most Na^+^-driven ATP synthase activity inferred by comparison to earlier bioinformatics-based estimations of Na^+^-binding protein ligands within ATP synthase subunits (109). Thus, Na^+^ driven ATP synthesis may be a common, albeit potentially non-critical characteristic among serpentinite-associated Bacteria and Archaea. Additional biochemical analyses are needed to better understand the selectivity of ATP synthase variants for Na^+^ or H^+^, how this may be leveraged to better predict selectivity based on sequence data (or structural prediction), and the level at which Na^+^ selectivity is needed for survival in serpentinization-influenced environments.

In addition to specific cellular adaptations to pH stress via additions and changes to cellular components, cells can also exhibit broader cell- or genome-wide adaptations to pH stress. For example, microorganisms have been hypothesized to mitigate environmental stress through genome streamlining (i.e., reducing genomic content via loss of unnecessary genes or genomic content), because genome replication is a particularly bioenergetically costly process (110). A common observation in multiple serpentinite-hosted communities is decreased average genome sizes in populations inhabiting environments with increasing serpentinization influence, as observed when comparing closely related taxa across pH gradients (39, 48, 53, 65) or when comparing genomes from whole communities across pH gradients (18, 52). Given the bioenergetic burden of growth at hyperalkaline pH combined with many serpentinization-associated anaerobic taxa likely operating metabolically near the thermodynamic limit of life, decreasing the energetic burden of genome replication via genomic streamlining may consequently represent a common adaptive strategy for cells in serpentinization-influenced habitats.

### 2.2 Inorganic Carbon and Adaptations to Overcome its Limited Availability

Alongside physico-chemical stress imposed by pH on cells, one of the most pressing issues for microorganisms inhabiting hyperalkaline waters is the lack of DIC for primary production. DIC availability in waters is inversely proportional to pH, owing to carbonate precipitation leading to very low DIC levels in many serpentinite systems (23, 42, 44, 49, 97). Consequently, a longstanding knowledge-gap in serpentinizing systems is understanding where the carbon that supports chemosynthetic biomass production is derived. Considerable evidence from ^14^C radiotracer and ^13^C uptake experiments (52) in conjunction with genomic evidence indicates the potential for carbon fixation of serpentinite populations through various canonical microbial pathways (27, 35, 52, 57, 111), although different carbon fixation pathways may be differentially distributed among communities based on the extent of serpentinization influence (65, 111). For example, the relative abundances of putatively autotrophic Bacteria and Archaea encoding WL pathways correlate with pH (i.e., serpentinization influence) in waters of the Samail Ophiolite (65). This is consistent with the WL pathway being one of the most energetically efficient carbon fixation pathways (11), likely helping to increase the fitness of cells experiencing stress associated with the reducing, hyperalkaline, and energy limited conditions of serpentinite systems.

Although DIC is limited at high pH, formate (HCOO^-^), a byproduct of serpentinization, can often be measured in many serpentinite systems (27, 41, 42, 112). Emerging evidence suggests that several putative autotrophs inhabiting the most hyperalkaline serpentinite environments may have adapted to chronic DIC limitation by favoring the utilization of HCOO^-^ as a carbon source. For example, genomic analyses of the “Type II” *Methanobacterium* spp. inhabiting the most hyperalkaline waters in the Samail Ophiolite indicate that they have restructured their core methanogenic pathways to potentially utilize HCOO^-^ as a source of both reductant (replacing H_2_-dependent [NiFe]-hydrogenases with HCOO^-^-dependent enzymes in the core methanogenesis pathway) and CO_2_ following its cytoplasmic oxidation by formate dehydrogenase enzymes (48). This is consistent with production of CH_4_ from ^14^C-labeled HCOO^-^ in microcosms containing Samail Ophiolite fracture waters where the Type II *Methanobacterium* were most abundant (48). More recent ^14^C conversion microcosm studies have shown production of CH_4_ from HCOO^-^ in rock-associated communities within microfractures of cored (50 m depth) peridotite rock from the Samail Ophiolite (56). Similar HCOO^-^-dependent adaptations to the core enzymatic steps of the WL pathway have been hypothesized for uncultured Acetothermia and Lithacetigenota bacteria from the Samail Ophiolite and the Cedars systems, respectively, based on genomic data (39, 65). Moreover, HCOO^-^ has also been implicated as a key carbon source in the LCHF and CROMO based on geochemical data and metagenomic analyses (111, 112). Perplexingly however, HCOO^-^ seems to not be used by the LCMS methanogens that dominate biofilms in chimneys but rather by lower abundance bacterial taxa present in the biofilms (36, 75). Intriguingly, nearly all of the above taxa, with the exception of some LCHF and CROMO bacteria (111), do not encode canonical HCOO^-^ transporters of Bacteria or Archaea, suggesting that yet to be discovered mechanisms of transport exist in these cells since HCOO^-^ is charged at pH >3.7 and consequently requires active transport into cells.

In addition to the above strategies, glycine reduction by H_2_ or formate to generate acetate has recently been hypothesized as a mechanism of biomass production in the Cedars based on metagenomic analyses (39). This could represent another potential energy conservation strategy that circumvents the need for DIC to drive primary production. Nevertheless, the glycine-based pathways, like many of the other pathways outlined above, remain hypothetical and the source of glycine (abiotic or biotic) to enable such a metabolism requires further investigation. As such, one of the most enigmatic questions of serpentinite ecosystem function – the source of carbon to support biomass production – remains unanswered. Genome-guided cultivation efforts or carefully designed experimental isotopic approaches (e.g., ^13^C-based isotope probing studies) could provide new avenues to identify the source of carbon that supports biomass synthesis in these systems and identify the members of communities responsible for such activities.

### 2.3. Hydrogen (H_2_) as a Primary Driver of Ecosystem Productivity

H_2_ is a product of serpentinization and, when accumulated to high concentrations, can drive the reduction of DIC to more reduced carbon species (5). Notably, CH_4_ is present at relatively high concentrations in serpentinites and is potentially formed through both abiotic and biotic mechanisms (44). As such, these sources of reductant (H_2_ and CH_4_) have been inferred to support communities hosted in serpentinites (113–115). Interestingly however, methanotrophs, apart from ANME in the LCHF (discussed above), appear to be generally limited in these environments, with variable evidence for their presence or methanotrophic activity in only a few systems like springs of the Voltri Massif (27) and Santa Elena ophiolite (116), despite that oxidants that can be coupled to CH_4_ oxidation (e.g., SO_4_ ^2-^) are generally present. Likewise, methanotrophic potential was only identified in less serpentinized (i.e., more oxidized) waters of the CROMO system (111), consistent with the necessity of coupling methane oxidation to oxidants like O_2_. Thus, methanotrophy does not appear to generally support primary productivity in most serpentinite systems.

In contrast, considerable evidence exists for H_2_ in supporting microbial metabolisms in serpentinites. Genes encoding the enzyme catalysts that allow for reversible H_2_ oxidation, of which there are three types categorized by their metallic cofactors ([FeFe]-, [NiFe]-, or [Fe]- hydrogenases) (104), are commonly identified among serpentinite populations (35, 41, 52, 117, 118) and are generally enriched among subsurface microbial populations (119). Several studies have indicated the importance and/or enrichment of hydrogenases among the genomes of microorganisms inhabiting serpentinite ecosystems. For example, elevated diversity and abundance of PCR-amplified hydrogenase genes were observed in diffuse venting fluids from the Quest site of the Logatchev deep sea hydrothermal field relative to fluids from a non- serpentinizing vent system (120). Likewise, a recent analysis of Samail Ophiolite fracture waters within boreholes indicated that they harbored communities where nearly 90% of populations (represented by MAGs) encode one or more isoforms of hydrogenase enzymes (20), far surpassing an estimate that 26% of archaeal and bacterial genomes encode hydrogenase homologs (104), indicating selection for hydrogenase-encoding populations within these waters. Moreover, the relative prevalence of hydrogenases among encoded proteins increased with increasing serpentinization influence (20), suggesting that cells in systems impacted most by serpentinization are adapted to capitalize on abundant H_2_.

To further examine this phenomenon, an analysis of hydrogenase homologs encoded in 68 publicly available serpentinite metagenomes (inclusive of the 70 described above but excluding two for methodological reasons; see Supplementary Methods for details) revealed a pH-dependent composition (**Fig. 3a**; *R*^2^=0.44, *p* < 0.001; envfit analysis), wherein the fractional abundance of hydrogenase homologs within metagenomes significantly increased with increasing pH (**Fig. 3b**, *R*^2^=0.31, p ≤ 0.001; linear regression; see Supplementary Methods for additional methodological information). Additional research is warranted to understand how the evolution of serpentinite-taxa and their associated hydrogenases, along with their coupling cofactors (e.g., NAD(H), Fd, F_420_, quinones, and cytochromes), has enabled various types of hydrogenotrophy and autotrophy in serpentinization-influenced ecosystems globally and throughout Earth history. Given that membrane-bound, extracellular [NiFe]-hydrogenases can be associated with accessory subunits that enable coupling to membrane-associated hydride carriers like quinones, thereby leading to translocation and accumulation of protons, an analysis of the prevalence of these hydrogenases is warranted in communities inhabiting the most serpentinized environments.

**Figure 3.**
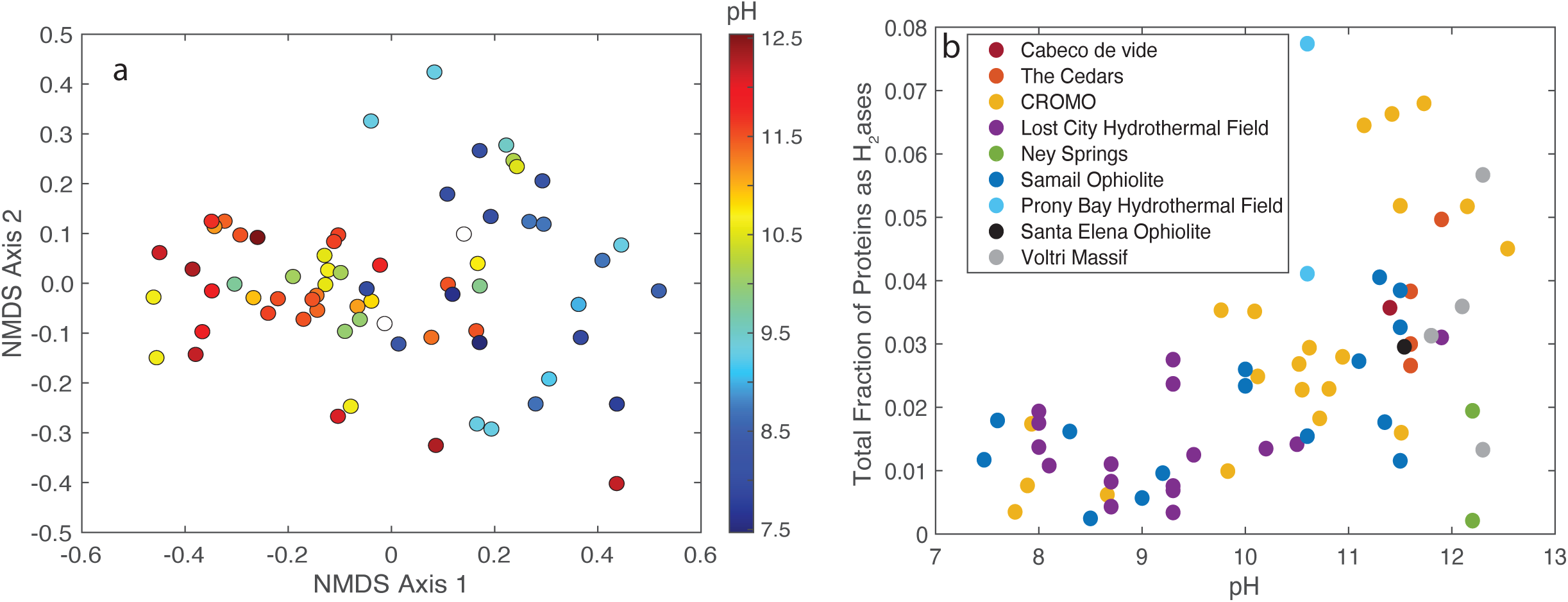
Variation in the composition and abundance of hydrogenase enzyme homologs encoded among 68 publicly available community metagenomes from serpentinization- influenced environments. **a)** Non-metric multidimensional scaling (NMDS) ordination of the hydrogenase profiles among 68 metagenomes, as predicted using the METABOLIC platform (described in Supplementary Methods). Points are colored according to the pH of the environment from where the metagenome derives, as indicated by the scale to the right. **b**) The total fraction of all hydrogenase orthologs relative to the total encoded proteins of a metagenome. Points are shown in relation to the pH of the environment from where the metagenome derives, with the system of origin indicated by color.

### 2.4 The Acquisition of Other Key Nutrients

Much of the attention on microorganisms inhabiting serpentinites has focused on mechanisms of energy conservation and carbon availability, with far less attention given to the availability and cycling of other critical nutrients needed to support productivity in serpentinite environments, including available nitrogen and phosphorous. The sources, sinks, and transformations of these elements remain enigmatic, although recent studies have provided some clarity. Within many systems, nitrogen substrates are not necessarily limiting. For example, organic nitrogen (e.g., in the form of amino acids) is particularly abundant in LCHF fluids, while inorganic forms of fixed nitrogen (e.g., NO_3_^-^ or NH_3_) are also available in biologically-relevant concentrations (121). Recent dual oxygen and nitrogen isotopic analysis of nitrogen species present in waters of the Samail Ophiolite suggested that nitrate (NO_3_^-^), which is typically detected in waters less influenced by serpentinization, likely derives from surface inputs and microbial nitrification. In contrast, isotopic evidence indicated that NH_3_ that is typically detected in waters more influenced by serpentinization, likely derives from microbial denitrification (122). Thus, *in situ* microbial nitrogen-cycling activities likely significantly control nitrogen availability in some systems. Nevertheless, both NO_3_^-^ and NH_3_ were generally detectable in Samail waters, indicating that fixed sources of nitrogen are not necessarily limiting in this ecosystem either. Moreover, the genomic capacity for nitrogen cycling via assimilation pathways, including di-nitrogen (N_2_) fixation (i.e., the microbial conversion of N_2_ to NH_3_), via nitrogenase can be abundant in the metagenomes and genomes from some serpentinite systems (24, 35, 122), altogether suggesting prevalent biogeochemical nitrogen cycling activities.

Among the organisms (assessed via MAGs) that are abundant in serpentinite systems and that encode the ancestral, molybdenum form of nitrogenase (Nif; (123)) are the putative sulfate reducers, *Thermodesulfovibrio* and *Desulfotomaculum* (26, 35), and the methanogen, *Methanobacterium* (48), all of which are likely to grow autotrophically using H_2_/CO_2_ or formate. The electron carriers (i.e., ferredoxin, Fd, flavodoxin) used by nitrogenases to reduce N_2_ to NH_3_ have low reduction potentials (*E^0^* ∼ -500 mV) that are lower than those of H_2_ or formate (*E^0^* ∼ - 500 mV). This thermodynamic issue is faced by autotrophic organisms (e.g., acetogens, sulfate reducers, and methanogens) that also rely on low potential Fd to drive the initial reduction of CO_2_ via the WL pathway (124). Many autotrophic organisms that fix N_2_ consequently bifurcate electrons from H_2_ (or formate) to drive the endergonic reduction of Fd to enable N_2_ reduction (125). The *E^0^* of H_2_ is strongly affected by pH and H_2_ concentration. Consequently, the reduction of low potential Fd in environments highly impacted by serpentinization (i.e., with hyperalkaline pH and high H_2_) has been suggested as favorable without bifurcation (70). A recent study demonstrated that the H_2_-dependent reduction of low potential Fd could be driven by native iron (Fe^0^) (126). Intriguingly, Fe^0^ (and native nickel, Ni^0^) minerals are associated with serpentinites Thus, it is possible that such native minerals (e.g., Fe^0^, Ni^0^, and awaruite (Ni_3_Fe)) in the presence of high concentrations of H_2_ could drive the abiotic reduction of N_2_, representing an alternate source of abiotic fixed N and a potential mineral precursor to nitrogenase enzymes Nevertheless, additional isotopic studies of nitrogen species in waters from natural serpentinites and laboratory studies of abiotic reduction of N_2_ by minerals commonly identified in serpentinite rocks are warranted to determine this possibility.

Perhaps even less understood than biological nitrogen cycling in serpentinites is how phosphorous is made available and cycled, given that phosphate is often present below the limits of detection (23, 43, 129, 130). A recent meta-analysis of 14 serpentinite-associated metagenomes suggested that the metabolic capacity to cleave phosphonates from organophosphates (e.g., methyl-phosphonates) via carbon-phosphorous lyase encoding genes were widespread and relatively abundant in serpentinite metagenomes (129). Nevertheless, the presence and provenance of such organo-phosphate compounds remains unknown in serpentinite systems and requires further study.

## 3. ARE SERPENTINITE BIOSPHERES SIMILARLY STRUCTURED?

### 3.1 Serpentinization as a Universal Control on Community Structures

Despite the variation in microbial communities among serpentinite systems described above, the common challenges that cells in these environments experience due to the process of serpentinization suggest that similar environmental factors likely shape the structure of serpentinite communities regardless of location. Consistent with this interpretation, microbial taxonomic diversity has been shown to be inversely correlated to pH based on 16S rRNA gene-based estimates of richness and diversity across Samail Ophiolite and CROMO subsurface waters exhibiting varying degrees of serpentinization influence (41, 42). Comparisons of the diversity in communities inhabiting highly serpentinization-influenced waters with those of nearby freshwaters (40, 47) or of those in waters less-influenced by serpentinization (46) are also consistent with these observations. Likewise, an analysis of the genomic diversity in the 70 publicly available metagenomes compiled for this review from nine globally distributed serpentinizing systems revealed similar patterns across the entire dataset (**Fig. 4a**; see Supplementary Methods for details). Specifically, pH was significantly inversely correlated (*R*^2^_adj_ = 0.25, *p* ≤ 0.0001) with the Nonpareil diversity metric, which is an assembly-independent estimate of genomic diversity based on sequence k-mer redundancy estimates (131). Thus, the environmental conditions that serpentinization generates limits microbial diversity and imposes strong ecological filters during the assembly of those communities. Nevertheless, significant geochemical variation both across and within serpentinites due to influences from processes in the deeper subsurface and those from the near-surface (e.g., mixing with seawater versus freshwater) likely further influence the composition of communities.

**Figure 4.**
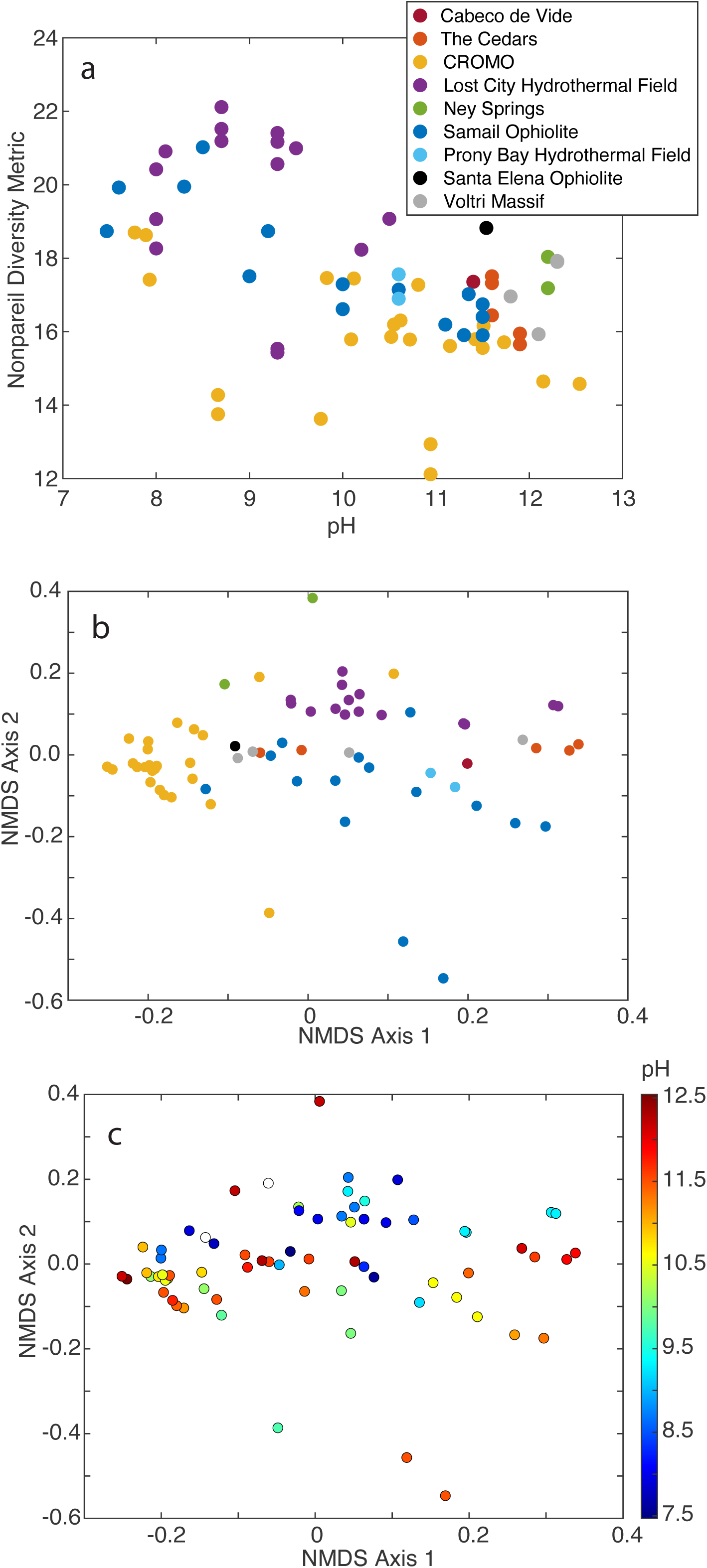
Metagenomic and taxonomic diversity of 70 publicly available community metagenomes from serpentinization-influenced environments. **a)** Metagenomic diversity, as measured by the Nonpareil diversity metric, plotted as a function of the pH of the environment where the metagenome derives (linear regression *R*^2^_adj_ = 0.25, *p* ≤ 0.0001). Points are colored by system of origin. **b** and **c)** Non-metric multidimensional scaling ordinations based on dissimilarity in the taxonomic profiles among communities, as evaluated with assembly-free analysis with the SingleM package (described in Supplementary Methods). Points are colored by their system of origin (**b**) as indicated in panel **a**, or by the pH of their sample based on the color scale to the right (**c**). Two samples did not have corresponding reported pH values and are shown as white circles. System of origin (**b**) was significantly associated with compositional variation based on envfit analysis (*R*^2^=0.56, *p* < 0.00), while pH (**c**) was only marginally associated with compositional variation (*R*^2^=0.09, *p* = 0.04). Additional sample information is provided in **Supplementary Table 1**.

Serpentinization is generally considered to occur in the subsurface of terrestrial and marine ultramafic rock-hosted systems. Deeply circulated waters impacted by extensive water/rock reaction processes (e.g., those with higher pH, higher gas concentrations, depletion of DIC, among other characteristics) can circulate back to the surface through networks of fissures or pore networks. Near-surface and oxidized freshwater or seawater can then potentially mix with the deeper fluids during their ascent to the surface, resulting in a continuum of serpentinization influence that is specific to a given system (15). Further, the specific geologic contexts of serpentinization-influenced waters impart additional influences on habitat characteristics. For example, waters venting at LCHF chimneys are highly variable in their chemical composition and are moderately hydrothermal, ranging up to 91°C, although the mixing of vented fluids with seawater can be substantial (112). In contrast, other recently studied marine serpentinite seep or chimney sites exhibit waters closer to ambient temperatures of < 10°C (22), while waters in continental serpentinite systems have temperatures largely near ambient or that are only slightly elevated (23, 41, 42).

The geochemical composition of vented or subsurface continental fluids largely reflects their sourcing from seawater (13), altered groundwaters (23, 42), or variable mixtures of the two in the case of near-coastal sites like the PBHF (New Caledonia) (132) and Ney Springs (California) (24). Despite the large potential for geochemical heterogeneity, many studies have documented that community composition is either qualitatively or quantitatively associated with proxies for serpentinization influence. Most commonly, pH is used as a proxy for serpentinization, although silica activity (which increases due to surface water influence) has also been widely used to infer the mixing of deeply sourced waters (133). Indeed, the degree of serpentinization is associated with community structural differences in systems where sampling across multiple site types occurs. For example, the contribution of deep vs. shallow groundwater was strongly correlated to community structure in the Cedars (47), as also observed in the Samail Ophiolite from 16S rRNA gene (42) and metagenomic-derived functional profiles (52). Likewise, the structure of communities from venting LCHF waters were chimney-specific, with a strong inferred influence of dynamic input of subsurface fluids, but weak inferred influence due to mixing with seawater (35). Similar interpretations have been suggested for the Old City Hydrothermal Field (22). Thus, a primary control on the microbial ecology of serpentinization systems is the degree of serpentinization influence on the associated waters.

Given that serpentinizing microbial ecosystems are largely driven by chemosynthetic communities whose metabolism is controlled by their specific geochemical environments, the null hypothesis for global serpentinization biosphere structure is that system-specific geochemical characteristics shape their overall structures. A recent meta-analysis of 16S rRNA gene profiles from four different serpentinite systems was consistent with this interpretation, with the microbial composition and structure of water samples primarily reflecting their system of origin (24). Likewise, an evaluation of the taxonomic composition of the 70 metagenomes compiled herein was consistent with this hypothesis, with system of origin strongly associated with overall taxonomic profile (envfit *R*^2^=0.56, *p* < 0.001; **Fig. 4b**), while pH (as a proxy for serpentinization influence) was minimally associated with overall composition (pH *R*^2^=0.09, *p* = 0.04) (**Fig. 4c**). Thus, localized conditions appear to largely dictate the composition of microbial taxa found in each system, despite that several individual taxa may be associated with globally distributed serpentinization systems, and that serpentinization influence structures communities within a given system, as described above and in **Section 4**. Unlike a recent meta-analysis of 14 serpentinite metagenomes, a distinction of LCHF community profiles relative to those from other systems was not apparent here (129), which could be due to the larger dataset size used here. Considering the metabolic functional potentials encoded by the above metagenomes (see Supplementary Methods for details), system of origin was again highly associated with functional variation (**Fig. 5a**; *R*^2^=0.55, *p* ≤ 0.001), while pH was considerably more associated with variation compared to the taxonomic profiles (**Fig. 5b**; *R*^2^=0.25, *p* ≤ 0.001). Thus, similar challenges imposed by serpentinization may lead to the convergence of functionalities (e.g., H_2_ metabolism as suggested by **Fig. 3**) among taxa that may be considerably differentiated taxonomically among systems, owing to localized characteristics.

**Figure 5.**
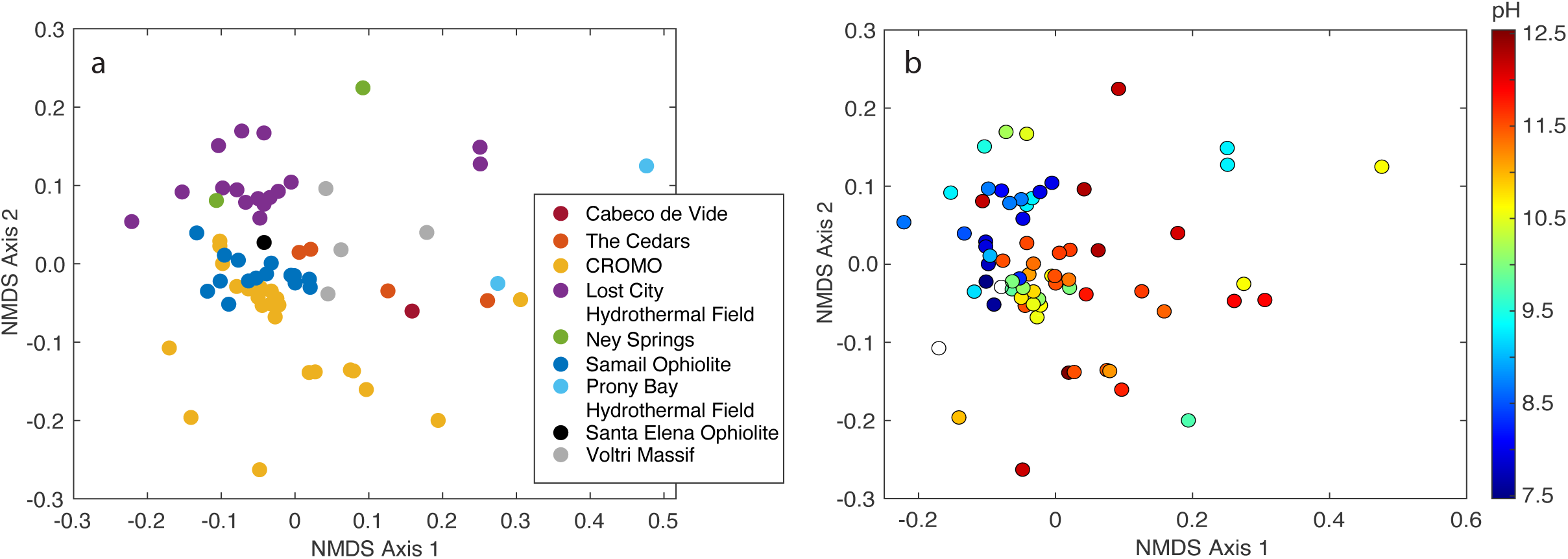
Variation in metabolic functional potential among 68 publicly available community metagenomes from serpentinization-influenced environments. **a)** Non-metric multidimensional scaling (NMDS) ordination of the functional potential profiles among 68 metagenomes (described in Supplementary Methods). Points are colored by (**a)** their system of origin, as indicated by the legend on the bottom right and (**b)** the pH of their sample, based on the color scale to the right. Two samples did not have corresponding reported pH values and are shown as white circles. System of origin **(a)** was significantly associated with compositional variation based on envfit analysis (*R*^2^=0.54, *p* ≤ 0.001), while pH **(b)** was also associated with compositional variation, but to a lesser extent (*R*^2^=0.25, *p* ≤ 0.001). Additional sample information is provided in **Supplementary Table 1**.

### 3.2 Within-system Heterogeneity

As suggested by the studies described above, the context of serpentinite systems influences the structure of their microbial communities and this occurs at various scales, including within the same system. The sharp redox gradients generated when reduced subsurface fluids mix with oxygenated fluids can generate distinct and extensive redox disequilibria. For example, redox gradients of >500 mV have been observed in only the top 40 cm of Cedars spring waters (47), while gradients of >750 mV have been measured along a 400 m interval in a recently drilled well in the Samail Ophiolite, with other wells similarly exhibiting extensive redox gradients (26). Recent studies have begun to explore depth-resolved differences of microbial communities in boreholes drilled at the CROMO and Samail Ophiolite sites. Analysis of waters across a gradient of O_2_ concentrations ranging from saturated at the surface to essentially 0% at ∼20 m depth at the CROMO site revealed contrasting geochemical and microbial community compositional gradients (134). Likewise, investigation across discrete, packed-off intervals along a 400 m depth profile in the Samail Ophiolite (to isolate individual fracture units) revealed discrete geochemical and microbial compositions that follow sharp redox gradients, with corresponding changes in the prevalence of aerobic and anaerobic taxa typical of serpentinite systems (59, 63). Although boreholes into actively serpentinizing systems for scientific investigation have only recently become available for study, they hold tremendous promise for understanding concerted microbial and geochemical dynamics as fluids from the deep subsurface and surface of serpentinizing systems mix. This is especially true for boreholes that intersect aquifers that harbor distinct water types and that could encompass the near total range of water chemistries found within a given system (26).

While many studies have examined spatial variation in serpentinite communities within or across systems, very little is known of their stability or temporal variation. A recent study of the Ney Spring system over a year revealed significant variation in the geochemistry of fluids that coincided with seasonal surface influences (e.g., air temperature, surface-derived biomass input) and substantial variation in “transient” low-abundance taxa (135). Nevertheless, the continuous presence of several taxa inferred to be residents of the deep subsurface serpentinized fluids sourcing the spring system was identified, suggesting a consistent supply of deep subsurface waters. Similarly, sampling of well waters in CROMO across multiple months within a year and across multiple years revealed that the assembly of those communities was largely structured by ecological filtering (i.e., deterministic factors) that was attributed to the significant ecological stressors inherent to serpentinite environments (136). However, transient changes in the communities were also observed, suggesting a secondary influence of stochastic processes (e.g., “drift” related factors). Analyses of the hydrology of the CROMO system revealed a general lack of communication between waters from different wells, which would provide a mechanism for ecological drift to occur if communities remain isolated for significant periods (136).

In the Samail Ophiolite, the deeply derived, highly serpentinization-influenced “Type II” waters have been estimated to have residence times of >20,000 years, with minimal communication with nearer surface, younger “Type I” waters (137). Analyses of mobile genetic elements and single nucleotide variants of closely related *Meiothermus* strains recovered from communities inhabiting Type I and II waters have suggested that long-term isolation of fluids, combined with limited gene flow among strains, promoted the diversification of these strains via parapatric speciation (63). Thus, the long-term stability of many serpentinite waters, combined with their unique geochemistry (relative to near surface waters), may provide a basis for ecological drift to drive microbial speciation. Additional temporal sampling of discrete fluid types, combined with isotopic and geochemical characterization to determine the extent of their isolation, could be used to begin to estimate the timeframes required for evolutionary change to occur in natural systems, in addition to population turnover times, of which very little is known in subsurface systems in general.

## 4. THE ORIGIN OF LIFE AND THE POTENTIAL FOR LIFE ON OTHER PLANETS

### 4.1 Do Serpentinite-Specific Taxa Exist?

The distinctiveness of microbial communities from highly serpentinized waters relative to less serpentinized waters or fresh-/marine-waters (e.g., (42, 47)) suggests that microbial taxa may be endemic to serpentinite ecosystems, despite the considerable variation in microbial composition across globally-distributed systems. The best evidence yet described for a serpentinization-specific lineage is the genus *Serpentinimonas* discussed above that is abundant and widespread in serpentinite environments (18, 41, 93), where it was first isolated. The lineage appears to be primarily associated with serpentinizing environments (18, 93), although it has also been identified in other hyperalkaline non-serpentinite systems like slag waste (138). The dependence of *Serpentinimonas* on H_2_-based autotrophy and their adaptation to hyperalkaline pH in contrast to their sister genus, *Hydrogenophaga* (18) suggests that *Serpentinimonas* may be specifically adapted to a lifestyle within serpentinization habitats.

Genome-resolved metagenomics approaches have also recently identified phylogenetically deep-branching bacterial lineages that are primarily associated with serpentinizing environments and that exhibit hypothesized metabolic adaptations to serpentinization-influenced environments as acetogen-like organisms. These include the order-level UBA7950 lineage of the Bipolaricaulota (previously ‘Acetothermia’) phylum (65) and the proposed ‘Lithacetigenota’ phylum (39). Members of the UBA7950 lineage are associated with hyperalkaline Type II waters of the Samail Ophiolite (65), and have also been identified in LCHF basement fluids (35). In addition, the organisms exhibit carbon acquisition and H_2_- dependent metabolic adaptations for highly serpentinized fluids that differentiate them from other Bipolaricaulota present in less serpentinized fluids (65). Notably, 16S rRNA gene studies of chimneys in the active serpentinization PBHF system also documented Bipolaricaulota / ‘Acetothermia’ (that phylogenetically affiliate with the UBA7950 group) as early chimney colonizers and potential members of basement fluids (35, 139). Likewise, *Ca.* ‘Lithacetigenota’ populations have been primarily identified in serpentinite environments and also exhibit carbon acquisition and H_2_-dependent metabolic adaptations for such environments (39). Alongside *Serpentinimonas*, these taxa represent some of the best evidence yet documented for serpentinite-specialized lineages, although further analyses of their global distributions and evolutionary ecology are needed to fully assess this hypothesis.

In contrast to the above, many of the taxa commonly associated with serpentinite environments exhibit broad environmental distributions, particularly within other subsurface environments, suggesting relatively recent adaptation to serpentinite environments, or a lack of specific adaptation to these environments. For example, *Methanobacterium* spp. and *Desulfotomaculum* spp. are commonly dominant in serpentinite communities but are also broadly distributed in other subsurface environments (140, 141), in addition to environments that are not hyperalkaline or subsurface-related (142, 143). Comparisons among serpentinization-influence gradients may provide critical insights into whether species or strains within these lineages exhibit specific adaptations to serpentinization habits. For example, the *Methanobacterium* Type II lineage described above exhibited hypothesized serpentinite-specific adaptations to mitigate limited carbon availability relative to closely related species found in less alkaline waters (48). Nevertheless, additional rigorous evolutionary analyses are needed to ascertain the extent to which taxa are specifically associated with serpentinites and the evolutionary scale at which this adaptation may have occurred (e.g., at the order/phylum level as in Acetothermia / “Lithoacetigenota” or at the genus/species level as in *Serpentinimonas*/Type II Methanobacteria.

### 4.2 Are Serpentinite Taxa Reflective of Those on Early Earth?

The simplicity of serpentinization, which requires only fractured ultramafic rock and circulating water to initiate the production of H_2_ and drive organic synthesis, has led to many serpentinization-related origin of life hypotheses on Earth and potentially, on other rocky planets (7, 9, 16, 115). Intriguingly, the collective chemical reactions in the abiotic synthesis of simple carbon compounds (e.g., CH_4_) from H_2_ and DIC are highly analogous to the stepwise organic carbon synthesis reactions in the autotrophic WL and core methanogenesis pathways of anaerobic Bacteria and/or Archaea described in above sections (16, 144). Consequently, the WL pathway has been long-associated with origin of-/early-life hypotheses in serpentinite systems. Recent studies of uncultured deep-branching acetogen-like organisms (39, 65–67) have provided new evidence for the WL pathway in serpentinite-associated lineages. Despite the prevalence of methanogens in serpentinite systems (that encode archaeal versions of WL pathways) (12), their relevance for understanding early archaeal evolution is less clear. For example, the methanogens or anaerobic methanotrophs that typically inhabit marine serpentinite systems (e.g., LCMS and ANME) belong to the Methanosarcinales and Syntrophoarchaeales orders that are considered recently-evolved (e.g., Class II) methanogens/methanotrophs (12, 145–147) (**Fig. 6**). In contrast, methanogens that typically inhabit terrestrial serpentinite systems, *Methanobacterium*, belong to the Methanobacteria class that are considered among the earlier- evolving (Class I) methanogens (12) (**Fig. 6**). However, the *Methanobacterium* most associated with highly serpentinization-influenced waters in the Samail Ophiolite appear to be derived from those inhabiting less-influenced waters and are nested well-within the Methanobacteria, suggesting that they are relatively evolutionarily recent (48). Likewise, no other methanogenic (or anaerobic methanotrophic/alkanotrophic Archaea) that are considered among the earliest to have evolved (12) have been identified in serpentinite systems. Many of the other bacterial or archaeal groups typically identified in contemporary serpentinizing systems including the Proteobacteria (e.g., *Serpentimonas*, *Hydrogenophaga*, other Burkholderiaceae), Nitrososphaeriales (ammonia oxidizing Archaea), Desulfurobacterota (previously Deltaproteobacteria sulfate-reducers) or Nitrospirota (*Thermodesulfovibrio*) are all considered later-evolving lineages (148–150) (**Fig. 7**). Thus, the extent that microbial phylogenetic evidence from contemporary serpentinite systems supports their reflection of early life on Earth is mixed.

**Figure 6.**
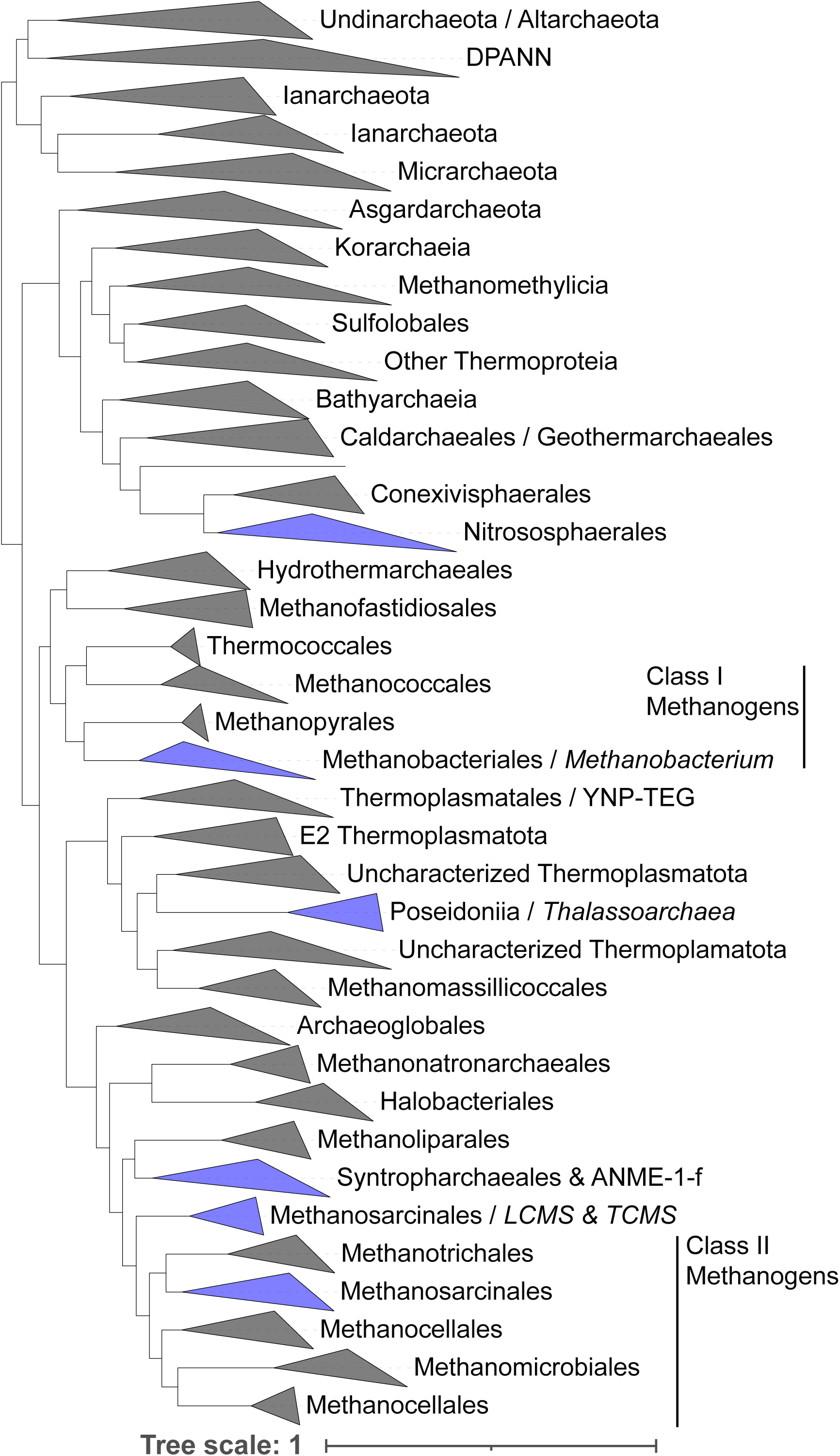
Phylogenetic reconstruction of major archaeal lineages highlighting those that are abundant in serpentinite systems. The tree was modified from the Maximum Likelihood phylogeny of archaeal orders described in Mei *et al.* 2023. The newick tree corresponding to Fig. 1a in the above study was used to highlight taxonomic lineages discussed in this review. Cohesive clades that coincided with taxonomic classifications are collapsed and shown as triangles, with those in blue corresponding to taxa that have been identified in serpentinite ecosystems and that are discussed in the main text. Notable taxa found in serpentinite communities are indicated after the forward slash and are indicated in italics. The tree was rooted with bacterial taxa (not shown). Scale shows expected number of substitutions / site.

**Figure 7.**
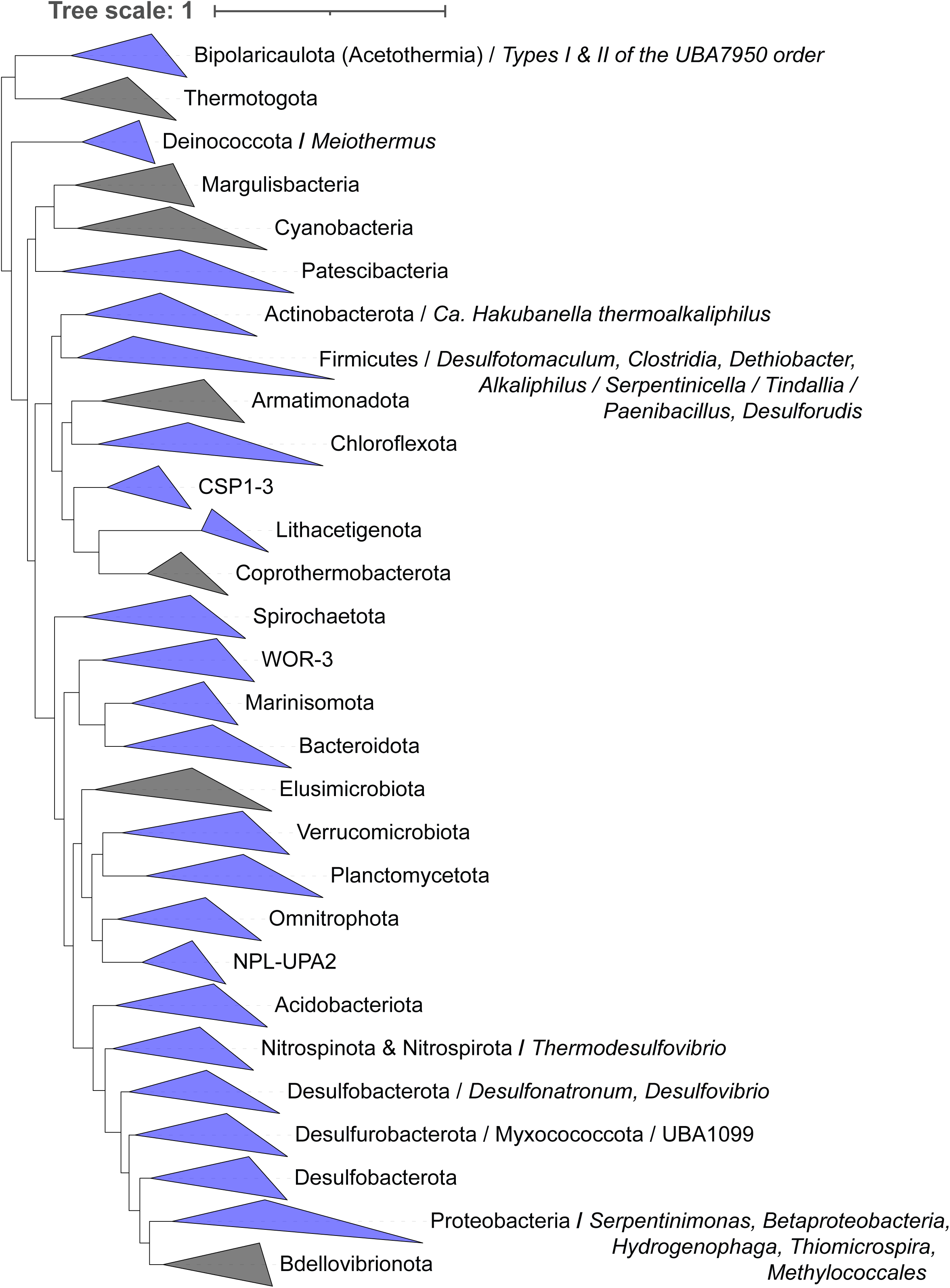
Phylogenetic reconstruction of major bacterial lineages highlighting those that are abundant in serpentinite systems. The tree was modified from the Maximum Likelihood phylogeny of bacterial orders described in Martinez-Gutierrez *et al.* 2021 (described in Supplementary Methods). Cohesive clades that coincided with taxonomic classifications are collapsed and shown as triangles, with those in blue corresponding to taxa that have been identified in serpentinite ecosystems and that are discussed in the main text. Notable taxa found in serpentinite communities are indicated after the forward slash and are indicated in italics. The tree was rooted with archaeal taxa (not shown). Scale shows expected number of substitutions / site.

The recently evolved nature of many of the microorganisms inhabiting modern serpentinites is likely attributable, at least in part, to the input of more oxidized near surface fluids today that overprint the reduced geochemistry and microbiology of those systems. On early Earth, some of the most readily available electron acceptors would have been CO_2_ and sulfite (SO_3_^2-^) (151, 152) that could support hydrogenotrophic acetogens/methanogens and sulfite (9, 152, 153). However, the progressive oxygenation of Earth over the past ∼2.3 Ga (154) allowed O_2_ and other high potential oxidants (e.g., NO_3_^-^, Fe(III), SO_4_^2-^) to become more available. The thermodynamics of H_2_ oxidation coupled to reduction of these higher potential oxidants (92) are more favorable for organisms that can respire them, allowing them to outcompete methanogens and acetogens for shared substrates like H_2_ or even CO_2_ (155). Alternatively, but not mutually exclusively, the cellular biochemistry of many anaerobes, including many methanogens and acetogens, is incompatible with O_2_ (156), further narrowing the extent of their niches. Likewise, Earth’s composite biosphere is far more productive today than it was prior to the origin of photosynthesis (157). Thus, it is likely that photosynthate infiltrates the near subsurface and can support a larger population of heterotrophic organisms in modern serpentinites than those on early Earth, in particular in highly productive tropical areas where serpentinites exist (e.g., as in the Zambales, Philippines (158)). While organic carbon can abiotically form and become stored in serpentinites (159), the restructuring of the global carbon cycle due to photosynthetically-derived carbon likely continues to profoundly affect contemporary serpentinite ecosystems. Finally, >3.8 Ga of evolution has improved the fitness of microorganisms through selective adaptations, with more fit populations displacing less-fit populations. Collectively, it is reasonable that the niches inhabited by acetogens, methanogens, and other anaerobes in serpentinites today are fundamentally different from and probably less extensive than they would have been on early Earth, again most notably in the near surface areas where mixing with oxidized fluids can more readily occur. Thus, future studies of the physiology, adaptations, and phylogenetic patterning of microorganisms across gradients of subsurface-surface influence are needed to help evaluate the extent that contemporary serpentinites and the communities they host reflect those of early Earth.

## 5. KEY OUTSTANDING QUESTIONS

The previous ∼20 years of study of serpentinite-hosted microbial ecosystems have fundamentally illuminated patterns in their diversity, function, and activity across geochemical gradients and their adaptations to conditions imparted by serpentinization. Nevertheless, several key outstanding questions regarding the microbial ecology of serpentinites remain unanswered. While common functional and ecological themes are observed across globally-distributed serpentinite systems, each system appears to host communities that are highly structured by nuances of their local environments. These observations point to unmeasured or yet to be considered geologic (bedrock mineralogy), hydrologic (fluid residence time, mixing), or chemical (organic carbon composition, source of nutrients) characteristics that may shape the microbial ecology of serpentinite systems. These observations also point to the need for continued exploration of both marine and terrestrial serpentinite systems to more fully explain emerging patterns in serpentinite microbial diversity, ecology, function, and activities. Further, while many key players in serpentinite-hosted microbial ecosystems have been identified, largely through cultivation-independent genomics studies, many of the mechanisms inferred to enable their habitation of those environments remain untested. For example, emerging evidence suggests that formate may be a key electron and carbon source in highly DIC-limited serpentinite systems, yet these assertions remain largely untested by laboratory or *in situ* studies. Thus, additional research is needed to identify the provenance of carbon that supports primary production, mechanisms of carbon acquisition/concentration, and *in situ* rates of activities. These insights will naturally lead to a better understanding of trophic relationships in these systems in addition to the extent of cell and nutrient turnover. Careful microcosm- and cultivation-based assays to better characterize activities under conditions that reflect *in situ* conditions will be key to furthering our understanding of these systems. For example, the rates of cellular turnover, the extent of cellular dormancy, and the tempo of microbial metabolism and nutrient cycling in these systems remains largely unknown. Experiments to address these areas will be challenging given that conditions in these systems abut the known limits of microbial habitability (e.g., lower limit of water stability, upper pH limit, lower energy limits), but such insights would be profound and will likely identify metabolic pathways and functions previously unknown in biology.

A key goal in the study of serpentinite ecosystems is understanding their relevance as analogs for studying origin of life processes on early Earth and the potential for life on other planets. However, many questions remain in these areas. Mixed phylogenetic support exists for contemporary serpentinite microbial taxa belonging to early-branching lineages that would reflect ancestry from earlier-evolving lineages, as would be expected if life originated and has been continuously maintained in these environments. However, nearly all contemporary environments considered as early Earth analogs that host early life (e.g., hydrothermal vents and some subsurface systems) are affected by overprinting by modern conditions due to the whole-scale co-evolution of Earth’s geosphere and biosphere over the last ∼3.8 Ga. Paramount among these is the evolution and proliferation of oxygenic photosynthesis and resulting increased biosphere productivity along with the average oxidation states of surface environments. Both changes would have eventually created new opportunities to challenge existing life in serpentinites. Further, many ecological niches would have collapsed due to competition or cells that were forced to retreat to deeper environments less influenced by near surface conditions and where their niches remained largely intact. Recent efforts to recover subsurface rock cores from the Samail Ophiolite as part of the Oman Drilling Project (160) hold promise in accessing subsurface niches that may be more reminiscent of early Earth. Nevertheless, additional efforts are needed to recover core (and fluids) from other global serpentinites. Until more comprehensive results of these efforts are available, care is warranted in extending observations from modern communities inhabiting serpentinites as analogs for those that would have been present on the anoxic early Earth or those that could be present on other rocky planets.

The process of serpentinization itself creates dynamic conditions that lead to changes in the chemistry and connectivity of subsurface fluids. For example, the volume of ultramafic rock can increase by upwards of 40% during hydration and alteration (161), creating stress that can be relieved through fracturing of rock. Further, mixing of reduced calcium-rich subsurface fluids with DIC-rich oxidized surface fluids can drive precipitation of calcite, both of which can alter fluid flow paths and connectivity. Notably, nearly all results from serpentinite systems involve the collection of waters (or sediments) from springs/vents or wells/boreholes within ultramafic rock settings. Natural serpentinite-hosted microbial populations most likely inhabit pores and microfractures that comprise highly segmented networks within rocks that are variably exposed to flowing water. When combined with the chemical evolution of serpentinite fluids, the potential for microbial populations to become spatially isolated in serpentinites over considerable time frames is extensive, consequently providing an impetus for incipient evolutionary divergence and speciation. Evidence in support of this hypothesis derives from a recent investigation of *Meiothermus*, which is a globally distributed and often abundant genus in serpentinites (49, 63) (Fig. 2). Specifically, two clades of an apparently Oman-specific *Meiothermus* lineage were identified in spatially isolated fracture waters, indicating a common ancestor. One clade was prominent in near surface, more oxidized aquifer waters and the other was more prevalent in deeper, more reduced aquifer waters. Analysis of single nucleotide variants and mobile genetic elements revealed detectable, albeit limited, evidence for gene flow/recombination among the populations. These observations were interpreted to indicate that that chemical variation generated by serpentinization, combined with physical barriers (reduced connectivity) that reduce/limit dispersal and gene flow, allowed for speciation of the *Meiothermus* populations. If similar studies could be conducted alongside accurate estimates of fluid exchange between aquifers, then it may become possible to quantify rates of gene flow and the time scales of *in situ* diversification among serpentinite-hosted taxa. Thus, coordinated sample collection and analyses between microbiologists and geochemists could leverage the unique properties of serpentinites to begin answering one of the greatest challenges in microbial ecology – quantifying rates of evolution in natural systems. Further, viruses are known to contribute to the tempo of microbial ecological and evolutionary processes in subsurface habitats (162). Yet almost nothing is known of viral abundances, diversity, effects on ecosystem processes, or effects on short-term evolutionary processes in serpentinite environments, representing a critical knowledge-gap that hinders a comprehensive understanding of the ecological and evolutionary tempo of serpentinite ecosystem processes.

Finally, nearly all understanding of the ecology of serpentinites derives from study of surface seeps/vents or fracture waters. Yet, in subsurface ecosystems, the bulk of microbial biomass is likely associated with surfaces as biofilms on minerals (14, 163, 164). Recent efforts to quantify biomass in microfractures in subsurface serpentinite rocks revealed six orders of magnitude variance, from 10 to 10^7^ cells/g in peridotite rocks from several >300 m cores from the Samail Ophiolite (26). Volumetrically, this range of cell densities bracket those for corresponding fracture waters in the Samail Ophiolite (52). Further, carbon cycling activities were shown to be ∼3 orders of magnitude higher (volumetrically) in rock microfractures than in corresponding bulk fracture waters, consistent with the notion that most of the biomass and thus biogeochemical activities of subsurface serpentinite ecosystems is associated with biofilms (56). Consistent with this hypothesis, H_2_ promoted carbon cycling activities of biofilms associated with peridotites from the equatorial Mid-Atlantic Ridge, with rock-hosted samples hosting greater biomass than overlying water samples (165). Nevertheless, little is known of the taxonomic and functional composition of serpentinite biofilm communities, despite that they may comprise the primary microbial consortia that influence biogeochemical cycling in serpentinite fracture networks. Much of this lack of information may arise from difficulty in recovering sufficient quantities of biomass or DNA from subsurface rocks, while concomitantly discounting potential contamination (as reviewed in (14)). Thus, technological advancements are needed to accurately evaluate the identity, characteristics, and activities of natural rock-hosted serpentinite communities. Significant and rapid advances are being made in computational capacity (e.g., through artificial intelligence and quantum computing), molecular methods (e.g., via ‘—omic technology scale and capacity), biochemical methods (e.g., in protein structure/function prediction), and spectroscopic analyses (e.g., in single cell activity measurements). Leveraging of these technological developments as they become available alongside carefully designed studies of serpentinite populations, communities, and ecosystems will certainly provide new insights beyond our initial insights gained in the last ∼20 years.

