## Supplementary Material for "The Microbial Ecology of Serpentinites"

**Supplementary Information**

**Summary**

Included in this file is a description of Supplementary Table 1 and Supplementary Methods that describe analytical details for the metagenomic analyses described in the main manuscript.

**Supplementary Table 1.** Sample and metadata information for the 70 serpentinite sample metagenomes analyzed in this review.

**Supplementary Methods**

**Compilation of Publicly Available Metagenomic Data**

Available metagenomic datasets were identified from previous studies of serpentinite microbial communities via literature and database searches. Only metagenomes that were publicly available in the National Center for Biotechnology Information (NCBI) Sequence Read Archive (SRA) were used, and in particular those that have been discussed in previous studies. Samples that clearly corresponded to biofilms were not considered. A total of 70 metagenome assemblies from water samples or substrate (e.g., chimney) composite samples were compiled (Supplementary Table 1). Given that available metadata (e.g., geochemical measurements) were highly sparse among the datasets derived from numerous studies, only sample pH was recorded, since it was the single geochemical parameter provided for all datasets. The raw reads were downloaded from the SRA and assembled, as described in detail elsewhere (1). Briefly, raw reads were trimmed of adapters and low-quality sequences using TrimGalore (v.0.6.6), down-sampled to remove sequence redundancy and improve subsequent assembly using bbnorm (bbmap v.38.7), and assembled using MetaSPADES (v.3.14.1). Total metagenome assemblies were subjected to protein annotation using PROKKA (v.1.4.6), considering only contigs of length ≥1,000 bp.

**Taxonomic, Diversity, and Functional Analyses**

Assembly-free taxonomic analysis was conducted using SingleM (v.0.18.0) using the ‘pipe’ function. Relative abundances of each taxon (identified by unique taxonomic classification strings) were estimated based on coverage values normalized to the total sum of coverage values for each metagenome. Only taxonomic classifications that were estimated to comprise >1% of the reads in any sample were used in subsequent visualizations and analyses. The relative abundances of the remaining taxonomic groups in metagenomes were then visualized using the ggplot2 package for R (v.4.4.1). The relative abundance matrix was also subjected to ordination analysis by transforming it into a dissimilarity matrix using the Bray-Curtis metric, followed by nonmetric multidimensional scaling via the metaMDS function of the vegan package for R (v.2.6-8) with 200 iterations. The analysis converged on a solution prior to 200 iterations, with a final stress 0.17, indicating reliable interpretability. The corresponding pH measurements for each sample and the system of origin were analyzed as correlates to the ordination configuration using the envfit function of vegan.

Metagenome reads were also subject to k-mer diversity analysis using Nonpareil (v.3.304) and with specification of a k-mer size of 19 that allowed comparable analyses among all metagenomes. The association of sample pH and Nonpareil diversity was evaluated using a linear regression in the base R package.

The metagenome-encoded protein annotations were subjected to functional analysis via the METABOLIC pipeline (v.4.0) and comparison against the Kyoto Encyclopedia of Genes and Genomes (KEGG) functional database, using a module presence cutoff of 0.75. All functions assigned by METABOLIC were used in the analyses to enable a comparison of overall functional variation among metagenomes. The number of hits for a given function within each metagenome were normalized to the total number of annotated functions for a metagenome to improve comparability across metagenomes that encompassed different sizes due to the use of different sequencing methods/strategies. The normalized matrix was subjected to NMDS ordination and fitting of pH/system of origin to the ordination, as described above for the taxonomic compositional analyses. To evaluate whether hydrogen gas (H_2_) metabolism was specifically functionally differentiated among systems or samples, the [NiFe]- and [FeFe]-hydrogenase functional annotations were analyzed separately, as described above for both the overall taxonomic and functional profile datasets. In addition, the association between hydrogenase homolog abundances (normalized to total proteins within a given metagenome) and pH were subjected to a linear regression, as described above.

**Phylogenetic Analyses**

To evaluate the phylogenetic distribution of major taxa present in serpentinite microbial communities, previously published, comprehensive archaeal (2) bacterial (3) phylogenies were used as frameworks. Specifically, the order-level phylogeny from Mei *et al.* 2023 (2) was retrieved via newick format and visualized in the interactive Tree of Life platform, with clades collapsed at the order level. In addition, the order-level (“balanced”) phylogeny within Martinez-Gutierrez et al. 2021 (3) was used as a framework to evaluate serpentinite bacterial phylogenies. Several notable lineages recently identified in serpentinite ecosystems were not present in the previously published bacterial phylogeny. Consequently, the genomes from the published order-level phylogeny were retrieved and combined with those from the candidate divisions “Acetothermia”/Bipolaricaulota, “Lithacetigenota”, and the NPL-UPA2 group that were recently identified or described in serpentinite systems (4-6). The genomes were collectively analyzed as previously described (1). Briefly, encoded proteins were annotated using PROKKA, and these were then subjected to searching for 30 housekeeping ribosomal proteins and RNA polymerase subunits using the markerfinder script (3). The resulting 30 protein datasets were individually aligned using clustalo, followed by concatenation of all 30 alignments into a super-matrix. The concatenated alignment was trimmed with trimal (specifying -gt 0.1) to remove low-quality alignment regions. The alignment was then subjected to Maximum Likelihood phylogenetic reconstruction using IQTREE (v.1.6.12), using the optimal substitution model chosen by the ‘TEST’ model finder algorithm (LG+I+G4), and 10 independent phylogenetic runs to compare analyses. 1,000 ultrafast bootstraps were used to evaluate node support.
